# No evidence for motor recovery-related cortical reorganization after stroke using resting-state fMRI

**DOI:** 10.1101/681320

**Authors:** Meret Branscheidt, Naveed Ejaz, Jing Xu, Mario Widmer, Michelle D. Harran, Juan Camillo Cortés, Tomoko Kitago, Pablo Celnik, Carlos Hernandez-Castillo, Jörn Diedrichsen, Andreas Luft, John W. Krakauer

## Abstract

Cortical reorganization has been suggested as mechanism for recovery after stroke. It has been proposed that a form of cortical reorganization (changes in functional connectivity between brain areas) can be assessed with resting-state fMRI. Here we report the largest longitudinal data-set in terms of overall sessions in 19 patients with subcortical stroke and 11 controls. Patients were imaged up to 5 times over one year. We found no evidence for post-stroke cortical reorganization despite substantial behavioral recovery. These results could be construed as questioning the value of resting-state imaging. Here we argue instead that they are consistent with other emerging reasons to challenge the idea of motor recovery-related cortical reorganization post-stroke when conceived as changes in connectivity between cortical areas.

## 1. Introduction

Spontaneous neurological recovery occurs in almost all stroke patients within the first months after the insult. While the underlying physiological changes that accompany spontaneous motor recovery in humans remain largely unknown, data from animal models have been interpreted as showing that cortical reorganization is a potential key mechanism mediating recovery (Dancause & Nudo, 2011; Grefkes & Ward, 2014; Nudo, 2006).

In the literature, the term cortical reorganization has been loosely defined and used to refer to any number of structural and physiological changes that follow injury. These changes can span the micro-, meso- and macro-scale, including synaptogenesis, axonal sprouting, and changes in cortical activation maps. We have argued elsewhere that the term functional reorganization should be reserved for those changes, including new cortico-cortical connections, that are causally related to or at least correlated with motor recovery (Krakauer & Carmichael, 2017). It should be added that reorganization has also been taken as a qualitative event, exemplified by the idea that one cortical area “takes over” another, which implies a change in the tuning of neurons, for example, when touching the face activates the hand area of sensory cortex in amputees. We argue elsewhere that a qualitative change in cortical representation need not be invoked to explain this result (Krakauer & Carmichael, 2017), but we will not use this definition here.

Evidence for functional reorganization after stroke comes primarily from studies of axonal sprouting. For example, Overman and colleagues (2012), in a mouse cortical stroke model, generated sprouting of axonal connections within ipsilesional motor, premotor and prefrontal areas by blocking of an axonal growth inhibitor (epinephrine A5). Similar results were reported for the neuronal growth factor GDF10 (Li *et al.*, 2015). Critically, however, in both studies no direct test of the relevance of axonal sprouting for motor improvement was performed, indeed not even a correlation with the degree of sprouting and behaviour was examined. In addition, most studies describing axonal sprouting after stroke found that it was cortico-subcortical instead of cortico-cortical connectivity changes that were linked to motor recovery (see e.g. Lee, 2004; Wahl *et al.*, 2014). Other studies that argue for a role of cortico-cortical connectivity changes underlying stroke recovery are limited by cross-sectional approaches or do not report behavior at all (Dancause *et al.*, 2005; Frost *et al.*, 2003; Liu & Rouiller, 1999; Napieralski *et al.*, 1996).

Despite the weak evidence for behaviorally-relevant new cortical connections in animal models post-stroke, these models have nevertheless led to widespread interest in identifying similar processes of functional reorganization in the human brain. One prominent non-invasive method is to measure inter-regional connectivity with resting-state fMRI (rs-fMRI; Biswal *et al.*, 1995; Fox & Raichle, 2007). This method relies on correlations between time-series of fMRI activity recorded while the subject is lying in the scanner without performing a task. Most often these correlations are computed between a set of pre-defined regions of interest (ROIs). The underlying assumption is that regions with connected neuronal processing show stronger statistical dependency of their spontaneous neuronal fluctuations. These correlations are commonly regarded as a measure of “functional connectivity”, which has been closely linked to structural connectivity (Friston, 2011; van der Heuvel *et al.*, 2009). In the context of stroke recovery, it has been suggested that reorganization can be detected as a change in such correlations/functional connectivity patterns (van Meer *et al.*, 2010). Specifically, for post-stroke recovery of hemiparesis, the advantage of task-free resting-state over task-based fMRI is that it avoids the performance confound (Krakauer, 2004, 2007); the connectivity measures are not biased by the inability of patients to match control performance due to motor impairment.

To date, results from rs-fMRI studies of functional connectivity changes after stroke have been mixed. Although, rs-fMRI studies have frequently found changes in interhemispheric connectivity patterns after stroke (Carter *et al.*, 2010; Chen & Schlaug, 2013; Golestani *et al.*, 2013), the direction of these changes and their correlations with behavior have been inconsistent. One study found a positive correlation between motor function and increased functional connectivity between the lesioned M1 and contralateral heterologous cortical areas (Park *et al.*, 2011), another study reported that interhemispheric homologous connectivity was associated with lower degrees of motor impairment but only for infratentorial strokes (Lee *et al.*, 2018). Yet another study showed that an increase in M1-M1 connectivity correlated negatively with motor function (Wang *et al.*, 2014).

There are many potential reasons for these inconsistencies in rs-fMRI findings. If patients with cortical lesions are included in the study design, it is possible to confuse changes in connectivity measures as a direct consequence of the lesion (e.g. the damaged area becomes disconnected from the brain) with changes associated with true reorganization. Additionally, most studies use different analysis protocols and measures to quantify changes in connectivity, making integration of evidence across studies difficult. Third, the majority of currently available studies have been cross-sectional but it is essential to evaluate changes in connectivity across the time-course of recovery.

To address these issues, we here report the results of a longitudinal rs-FMRI study of stroke recovery in patients with hemiparesis after subcortical stroke. Only patients with subcortical lesion locations were included in this study so that any changes in cortical connectivity could not be attributed to the presence of the lesion itself. We provide a detailed characterization of inter- and intrahemispheric connectivity between five cortical motor areas. Because of considerable variation of analysis approaches in the existing literature, in addition to our primary analysis, we also compared results after using two different pre-processing procedures, report results from an individual M1-M1 ROI analysis, and replicated the analysis approach from the largest longitudinal resting-state stroke study published to date (Golestani *et al.*, 2013).

## 2. Results

The main goal of this study was to determine whether motor impairment recovery following stroke was associated with systematic changes in cortical connectivity. Our two main questions were: 1) Is there a mean difference in the connectivity pattern between five motor regions (S1, M1, PMv, PMd, SMA) when comparing patients and age-matched controls at any time-point during stroke recovery? 2) Is there a change in patients’ connectivity patterns over time that is related to motor impairment? We analyzed data from 19 patients with subcortical stroke and 11 healthy controls. Behavioral assessments and resting-state images were obtained at five different time-points over one year. Each patient completed on average 4.5 ±0.7 sessions, with the overall experimental data being 89.5% complete (see also Table S1 for demographics and completed sessions in the supplemental material). We begin by quantifying the extent of impairment and recovery of upper extremity deficits in our patients in the year following stroke.

### 2.1. Patients showed substantial clinical recovery after stroke

We measured initial impairment and subsequent recovery of the upper extremity using the upper extremity portion of the Fugl-Meyer score (FM-UE), the Action Research Arm Test (ARAT), and hand strength (Xu *et al.*, 2017).

At the acute stage, all behavioral measures indicated impairment of the upper extremity for patients relative to controls (FM-UE: t(28)=3.706, p=0.001, ARAT: t(28)=2.315, p=0.028, strength: t(28)=5.195, p<0.001, Figure 1). These deficits recovered substantially over the course of one year, with the largest changes observed within the first three months (Week effect for FM-UE: χ^2^=24.865, p<0.001; ARAT: χ^2^=13.942 p=0.007; hand strength: χ^2^=13.419, p=0.009). No significant changes were observed in controls for any of the three measures.

**Figure 1:**
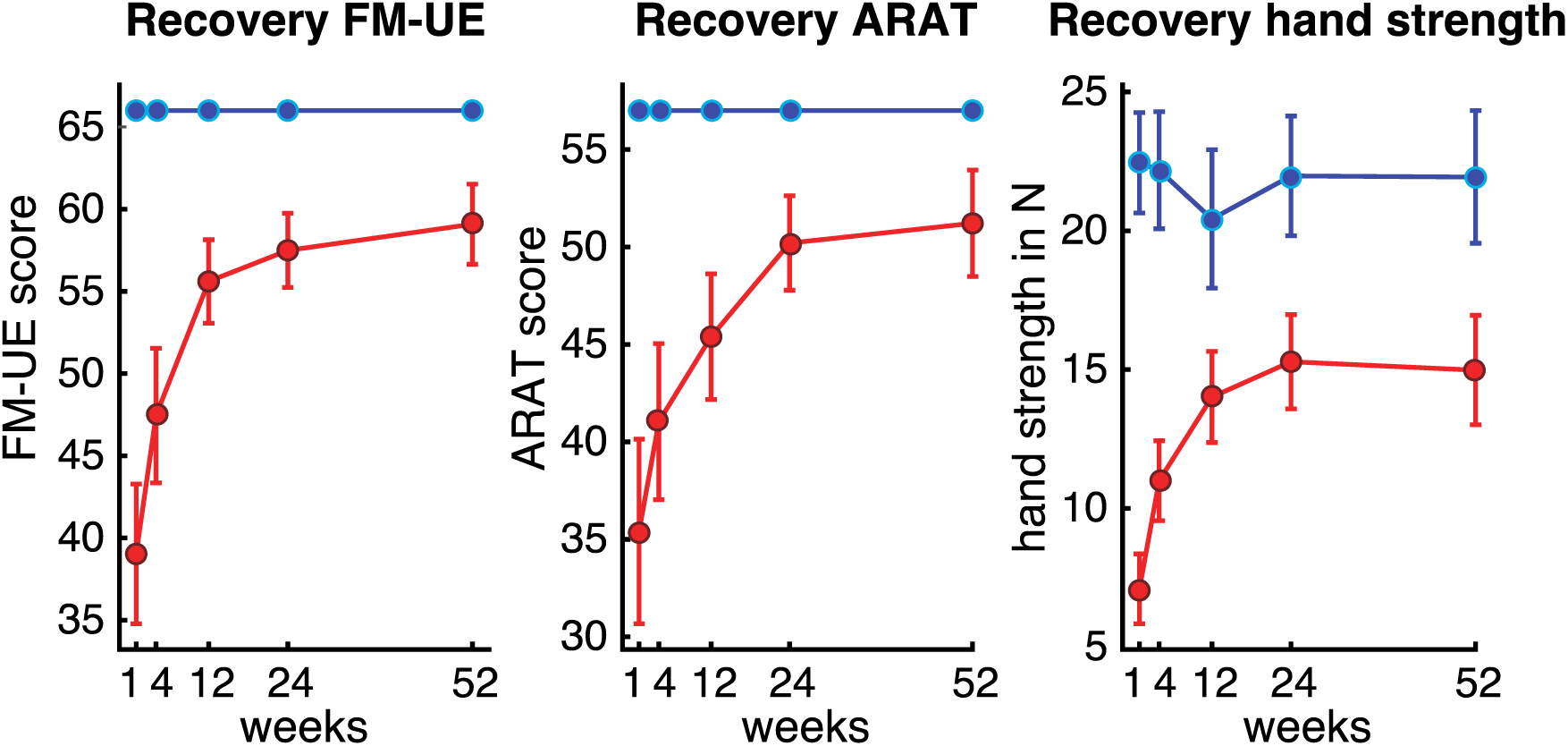
Recovery of upper extremity deficits after stroke over one year. For all behavioral assessments, the largest changes in recovery were seen within the first three months. Patients reached a plateau at 6 months and, on average, remained impaired compared to controls at all time-points. Note that patients had moderate to severe upper extremity impairment in the acute stage. Red lines = patients, blue lines = controls, FM-UE = Fugl-Meyer score Upper Extremity, ARAT = Arm Research Action Test.

### 2.2. Connectivity patterns across sensorimotor areas were reliable and stable in controls

Next, we looked at changes in connectivity patterns (pattern of ROI-ROI connectivity weights) between five key sensorimotor areas to determine if and how connectivity between these sensorimotor areas changed alongside behavior during recovery. To determine the connectivity patterns, we calculated pairwise correlations between the averaged time-series of BOLD activities between all possible ROI pairs to get a 10×10 matrix of connectivity weights (see Methods). An average connectivity pattern for patients and controls is shown in Figure 2a.

**Figure 2:**
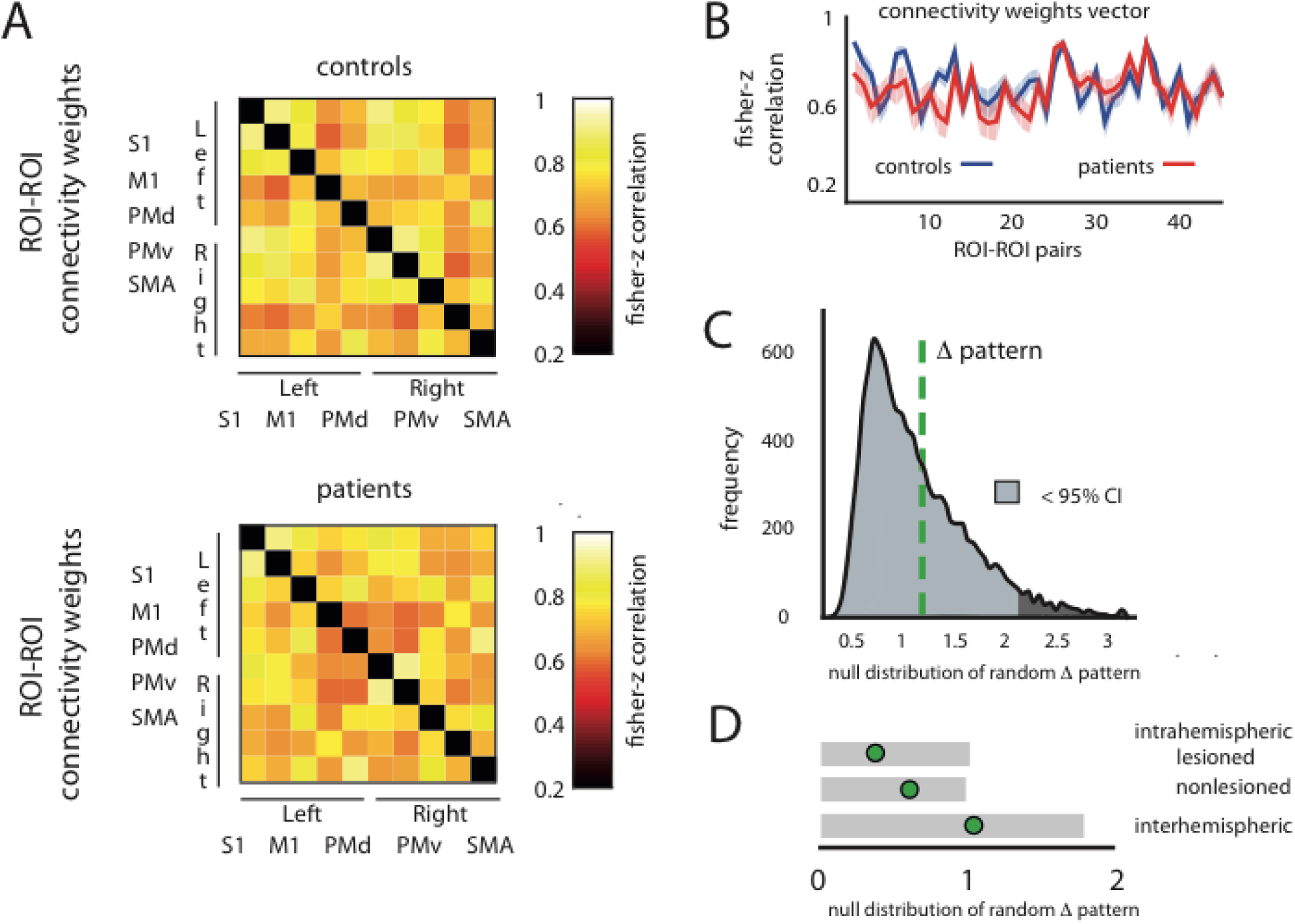
No systematic differences in connectivity patterns of patients and controls in the acute recovery period (week 1-2). (**A**) Heat map representation of average connectivity weights for controls and patients at the acute stage after stroke. The y- and x-axis show the five ROIs (S1, M1, Pmd, PMv, SMA) for the left and right hemisphere creating a connectivity matrix. One small square represents the connectivity weight for the respective ROI pairing. The diagonal (black) is missing, as it is the correlation of a ROI with itself. (**B**) Vectorized upper triangular part of the correlation matrix for the average full connectivity pattern of controls (blue line) and patients (red line). (**C**) To quantify the differences between connectivity patterns for controls and patients, we calculated the Euclidian distance between the two pattern vectors (Δpattern, dashed green line). The Euclidian distance is sensitive to differences of shape and scaling of patterns. The measured distance was then tested against the expected distribution if there were no differences between the two groups. To generate an empirical estimate of this distribution, we randomly shuffled group assignments and repeatedly computed the Euclidean distance (x10.000 times, histogram with frequency on the y-axis and absolute value of the Euclidian distance on the x-axis). For the acute stage after stroke, the Δpattern lay within the lower 95% percentile (grey shaded area) of the null-distribution. (**D**) The measured Δpatterns (green circle) for the intrahemispheric lesioned, non-lesioned or interhemispheric ROIs also always fell within the lower 95% range (grey boxes).

Connectivity patterns were highly reliable for both groups with high intrasession reliabilities (*all connections*, controls: R=0.66, CI 0.62–0.71, patients: R=0.70, CI 0.66–0.74; see supplementary material for inter- and intrahemispheric connections, Figure S2). An unbalanced mixed-effects ANOVA (see Methods) showed that the intrasession reliability was not significantly different between groups (χ^2^(1)=1.0782, p=0.2991) and showed no changes over time (controls: χ^2^(4)=6.174, p=0.187; patients: χ^2^(4)=1.922, p=0.75).

Furthermore, connectivity patterns for controls were stable, showing no significant change over time (*all connections:* Δweek acute_W4=0.841, confidence interval (CI) 0.597– 2.727; acute_W12=0.689, CI 0.582–2.412; acute_W24=1.079, CI 0.687–2.821; acute_W52=1.059, CI 0.611–2.531). Thus, for all subsequent analyses connectivity patterns for controls were averaged over time-points.

We also confirmed that the connectivity pattern for controls reflected known anatomical connectivity (Damoiseaux & Greicius, 2009). Within one hemisphere, the highest correlations were found between S1-M1 (0.91 ±0.47, Fisher-Z transformed), while the weakest correlation was found between M1-PMv (0.58 ±0.39). Between hemispheres, S1_right_-S1_left_ demonstrated the highest correlation (0.9 ±0.43), while M1_right_-PmV_left_ showed a weaker correlation (0.59 ±0.37). For correlations between hemispheres, homologous ROIs (e.g. M1-M1 or S1-S1) showed higher correlations of the BOLD time series compared to heterologous ROI-ROI connectivity weights (e.g. M1_right_-Pmv_left_ or S1_left_-Pmd_right_) as expected from interhemispheric neural-recordings (Asanuma & Okamoto, 1959, see supplemental results and Figure S3 for comparison of homologous versus heterologous interhemispheric connectivity).

### 2.3. There were no systematic changes in connectivity patterns in the acute recovery period

If disruption of the cortical projections through subcortical stroke leads to an acute reorganization of cortical circuits, one would expect that (on average) acute connectivity patterns of patients and controls would be different. Connectivity patterns for patients and controls were highly correlated in the early period after stroke (acute stage: R=0.69, p=0.0002; see connectivity matrices in Figure 2A and also Figure S4). To statistically test for significant differences between connectivity patterns, we used the Euclidian distance between the two groups’ mean patterns and compared it to a null-distribution obtained by a permutation test (Figure 2c). We found no systematic difference between patients and controls at the acute stage (Δpattern=1.246, CI 0.575–2.467). This was also true when only considering intrahemispheric connections of either the lesioned (Δpattern=0.367, CI 0.205–1.109) or non-lesioned side (Δpattern=0.603, CI 0.196–1.1) or interhemispheric connections (Δpattern=1.027, CI 0.394–1.968).

Even though the averaged connectivity patterns for patients and controls were indistinguishable at the acute stage, the heterogeneity in lesion locations for different patients might result in idiosyncratic shifts in connectivity patterns that in the whole group would be reflected as higher variability in patterns. To measure this within-group variability, we calculated the average Euclidian distance of each patient’s pattern to the patient group mean pattern and did likewise for controls. The average within-patient distance was 2.955, whereas the average within-control distance was 2.813, resulting in a difference of 0.142 (Δvariability). We compared this value to a null distribution of Δvariability generated with permutation testing. We found that resting-state connectivity patterns of patients showed a higher idiosyncratic, non-systematic variability compared to controls: The difference between the variability lay outside the 2.5% – 97.5% confidence interval generated by permutation testing (CI 0.018–0.051, Figure 3). Note that the confidence interval was not symmetric around zero, as the N for controls was smaller than for patients.

**Figure 3:**
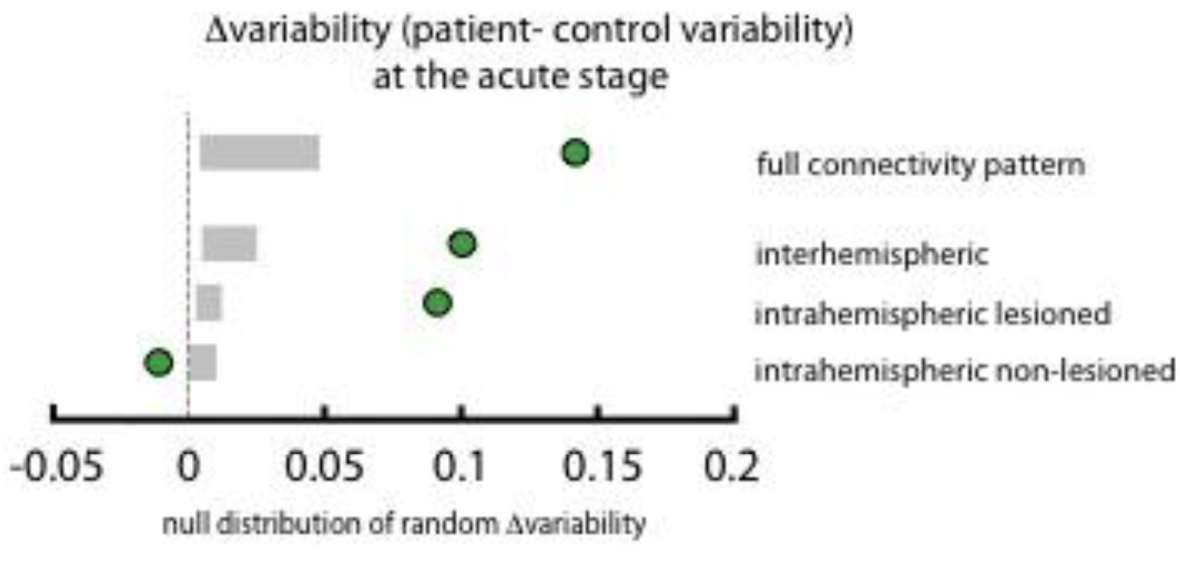
Patients showed a higher unsystematic variability compared to controls at the acute stage (**Δ**variability = green circle, 2.5%-97.5% range = grey boxes). Only for intrahemispheric non-lesioned ROI’s patients showed a lower variability.

The difference in variability for intrahemispheric lesioned and interhemispheric connections was also higher for patients. For intrahemispheric non-lesioned connections, we found higher variability in controls (intrahemispheric lesioned: Δvariability=0.091, CI 0.002– 0.023; non-lesioned: Δvariability=-0.01, CI -0.003–0.015; interhemispheric: Δvariability=0.1, CI 0.008–0.05, Figure 3).

Thus, overall, while connectivity patterns for patients were more variable, patient connectivity patterns were indistinguishable from control patterns at the acute stage.

### 2.4. There were no changes in patients’ connectivity patterns over time

Even though there were no systematic differences between connectivity patterns of patients and controls at the acute stage, we might expect to find changes in patient connectivity patterns over time as they recover from impairment.

We therefore quantified Euclidean distances between the average connectivity patterns at the acute stage as reference versus all other weeks (Δweek). Surprisingly, patients showed no increase in Euclidian distances between the acute stage and consecutive weeks (Figure 4 and Table 1).

**Figure 4:**
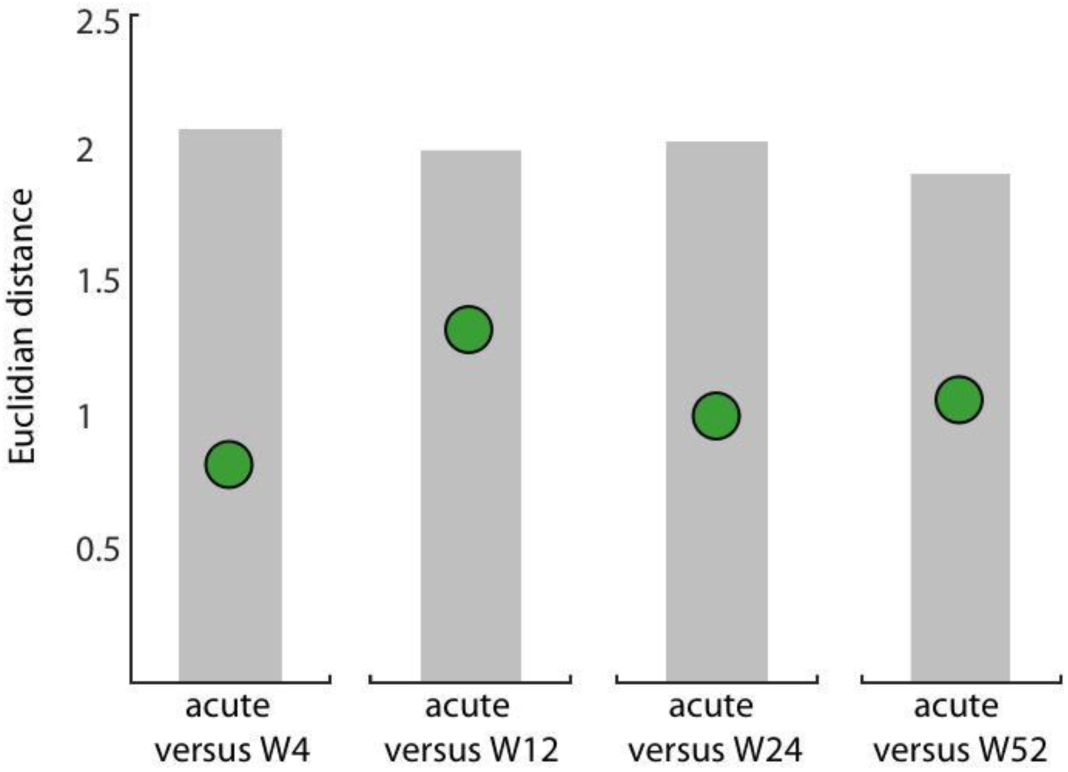
No significant change from patients’ acute connectivity pattern compared to time-points at the subacute or chronic stage. We computed the Euclidian distance between the average connectivity pattern of patients at the acute stage and all consecutive weeks (W4, W12, W24, W52; Δweek = green circles). Range of the expected distribution if there were no differences between the two groups (grey shaded area).

**Table 1:** Euclidian distances between the connectivity pattern of the acute stage compared to all subsequent time-points in patients for only interhemispheric, intrahemispheric lesioned, or non-lesioned subsets.

| Patients | acute_W4 | acute_W12 | acute_W24 | acute_W52 |
| --- | --- | --- | --- | --- |
| All connections | Δweek: 0.814,<br>(0.467 - 2.066) | Δweek: 1.322,<br>(0.488 - 1.994) | Δweek: 0.994,<br>(0.471 - 2.024) | Δweek: 1.063,<br>(0.512 - 1.898) |
| Interhemispheric | Δweek: 0.636<br>(0.309 - 1.695) | Δweek: 1.018<br>(0.3255 - 1.627) | Δweek: 0.665<br>(0.315 - 1.589) | Δweek: 0.77<br>(0.343 - 1.616) |
| Intrahemispheric<br>lesioned | Δweek: 0.353<br>(0.241 - 1.246) | Δweek: 0.413<br>(0.237 - 1.169) | Δweek: 0.457<br>(0.252 - 1.324) | Δweek: 0.558<br>(0.274 - 1.158) |
| Intrahemispheric non-lesioned | Δweek: 0.367<br>(0.152 - 0.859) | Δweek: 0.735<br>(0.157 - 0.886) | Δweek: 0.579<br>(0.158 - 0.919) | Δweek: 0.474<br>(0.174 - 0.805) |

As it could be expected from these results, patients showed reliably high correlations of their connectivity patterns with controls at the subacute or chronic stage (W4: R=0.74, p<0.0001; W12: R=0.76, p<0.0001; W24: R=0.87, p<0.0001; W52: R=0.80, p<0.0001) and no significant difference to control patterns 9 (Table 2). The analyses for intra- or interhemispheric connections alone found the same result (Table 1 & 2 and Figure S4).

**Table 2:**
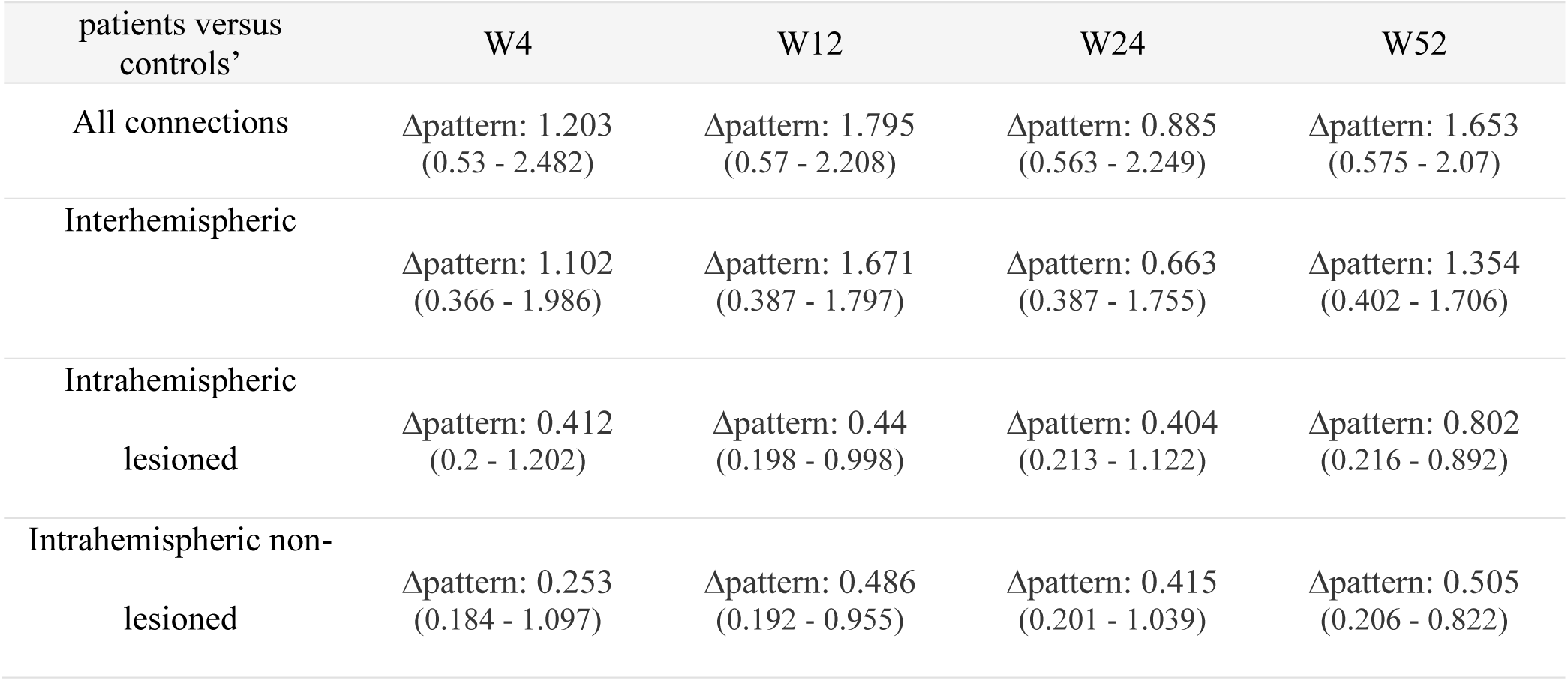
Difference between the connectivity pattern of patients compared to controls at Week 4, Week 12, Week 24, and Week 52 for only interhemispheric, intrahemispheric lesioned, or non-lesioned subsets.

| patients versus controls' | W4 | W12 | W24 | W52 |
| --- | --- | --- | --- | --- |
| All connections | Δpattern: 1.203<br>(0.53 - 2.482) | Δpattern: 1.795<br>(0.57 - 2.208) | Δpattern: 0.885<br>(0.563 - 2.249) | Δpattern: 1.653<br>(0.575 - 2.07) |
| Interhemispheric | Δpattern: 1.102<br>(0.366 - 1.986) | Δpattern: 1.671<br>(0.387 - 1.797) | Δpattern: 0.663<br>(0.387 - 1.755) | Δpattern: 1.354<br>(0.402 - 1.706) |
| Intrahemispheric<br>lesioned | Δpattern: 0.412<br>(0.2 - 1.202) | Δpattern: 0.44<br>(0.198 - 0.998) | Δpattern: 0.404<br>(0.213 - 1.122) | Δpattern: 0.802<br>(0.216 - 0.892) |
| Intrahemispheric non-lesioned | Δpattern: 0.253<br>(0.184 - 1.097) | Δpattern: 0.486<br>(0.192 - 0.955) | Δpattern: 0.415<br>(0.201 - 1.039) | Δpattern: 0.505<br>(0.206 - 0.822) |

By examining Euclidian distances between the individual connectivity patterns to the average connectivity pattern, we found a greater non-systematic variability in patients than in controls at the acute stage. However, the idiosyncratic variability of patients themselves did not change from the acute stage compared to the following time-points (Table 3).

**Table 3:**
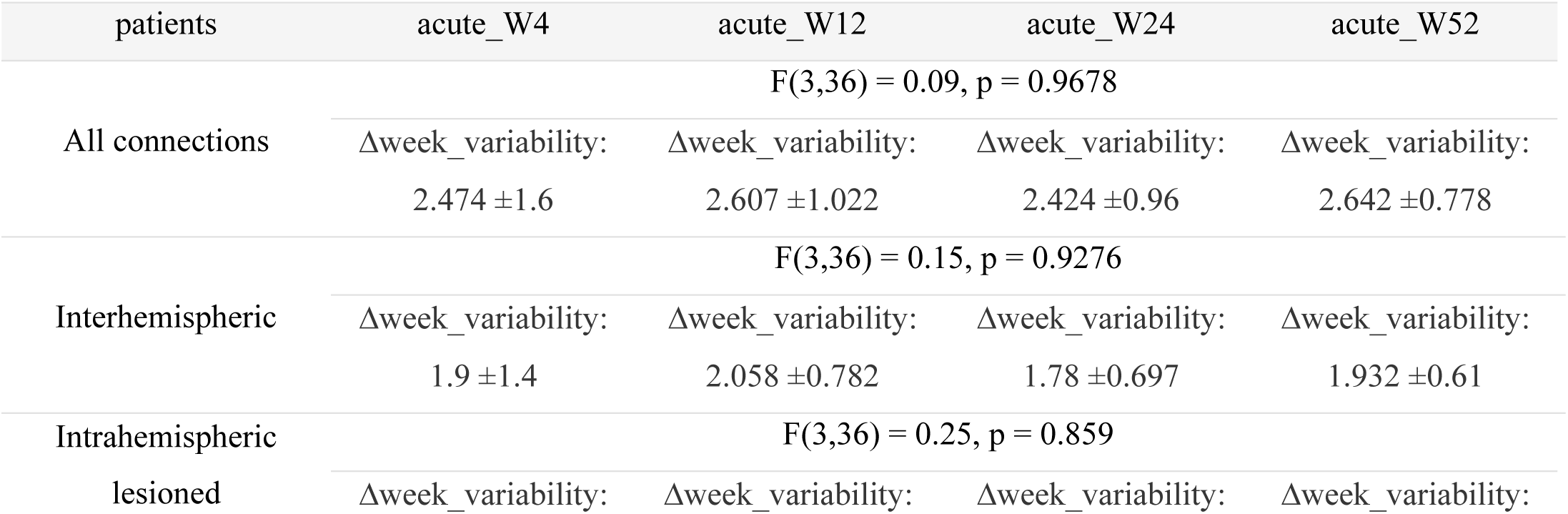
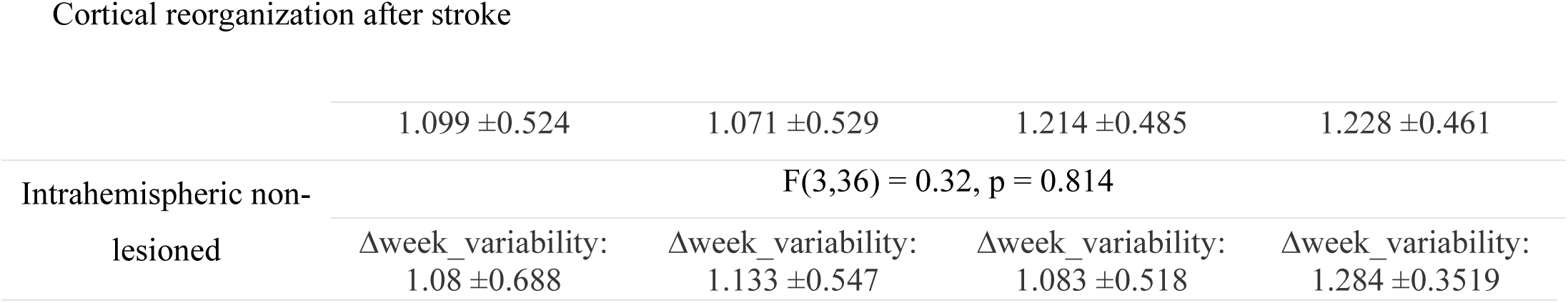
Difference in connectivity pattern variability in patients over time for all connections, interhemispheric, intrahemispheric lesioned, or non-lesioned subsets.

In summary, we found no evidence for a mean difference of connectivity patterns between patients within one year. More importantly, patients did not show any significant longitudinal change in connectivity patterns either systematically or regarding their group variability.

### 2.5. Comparison between alternative metrics for M1-M1 connectivity

Above we looked at the entire connectivity pattern between five sensorimotor areas within and across hemispheres and found no changes for patients either longitudinally or when compared to controls. In contrast, some previous studies have focused on individual ROI-to-ROI connections and have reported changes after stroke (Thiel & Vahdat, 2015). Specifically, changes in interhemispheric connectivity between the two motor cortices have been frequently reported (Carter *et al.*, 2010; Chen & Schlaug, 2013; Golestani *et al.*, 2013; Park *et al.*, 2011).

To test this finding, we investigated changes of interhemispheric M1-M1 connectivity weights over time and between patients and controls in our data set. The analysis showed a significant difference between patients and controls, with patients having a slightly lower average correlation between motor cortices (Figure 5a; *mixed model, group effect:* χ^2^(1)=5.759, p=0.016). Congruent with our other results, however, we found no longitudinal changes either for patients (*patient<u>_</u>week:* χ^2^(4)=5.836, p=0.212) or controls (*control_week:* χ^2^(4)=0.4.723, p=0.317).

**Figure 5:**
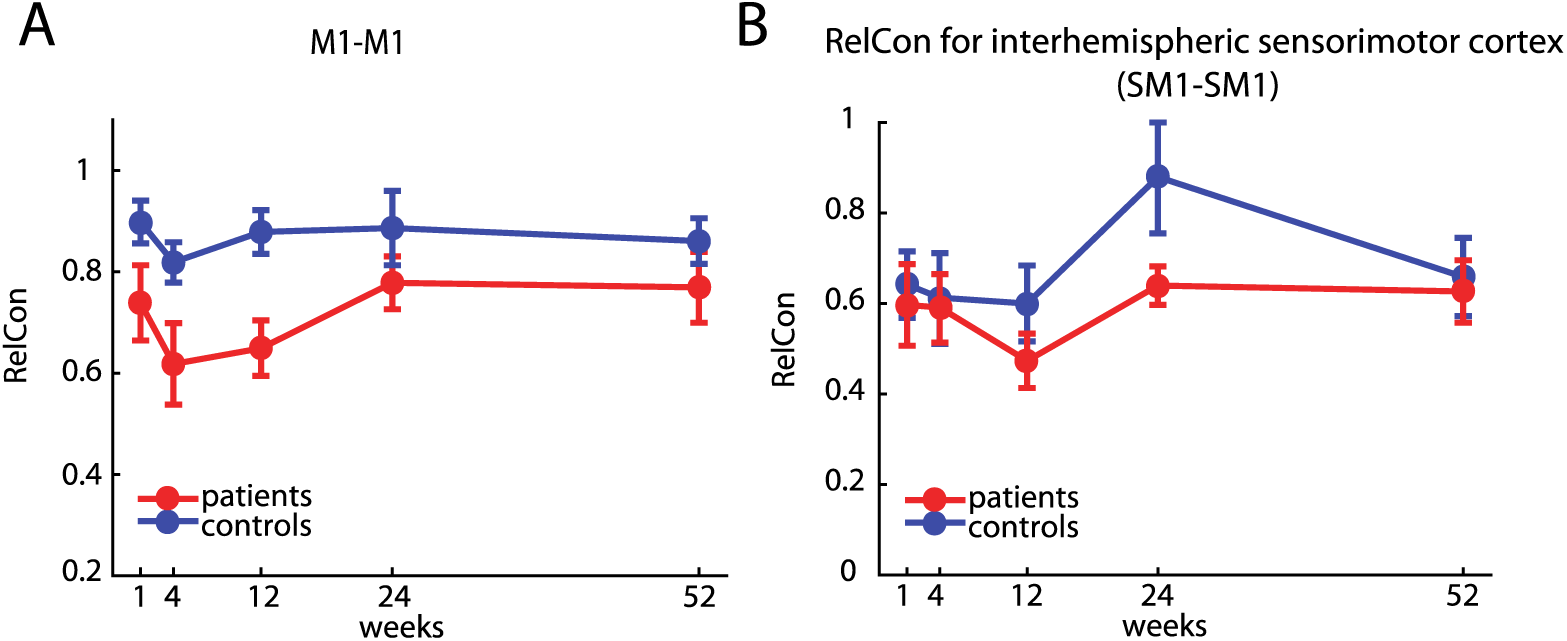
A) M1-M1 connectivity in our dataset. In patients, interhemispheric connectivity between the two motor cortices was systematically lower than compared to controls at all time-points. However, no changes of M1-M1 connectivity over time were found. B) Relative Connectivity of SM1-SM1 in controls and patients. While there was a significant difference in SM1-SM1 connectivity between the two groups, with lower RelCon for patients, there was no significant change over time.

Our results also contrast with another published finding that used an alternative metric of connectivity to assess changes in functional connectivity after stroke. Golestani and colleagues (2013) used a relative connectivity (RelCon, see Methods) measure between the two sensory-motor cortices and reported lower relative interhemispheric sensorimotor (SM1 RelCon) connectivity in stroke patients with a motor deficit compared to controls and stroke patients without impaired motor function.

Similarly, our patients had a lower RelCon for SM1-SM1 compared to controls at all time-points. Using a mixed-model, we found a significant difference between the groups (χ^2^(1)=5.2457, p=0.022). However, consistent with our results reported above, we did not find a change over time for RelCon SM1-SM1 in neither controls (χ^2^(4)=2.8087, p=0.5903) nor in patients (χ^2^(4)=8.2243, p=0.0837; Figure 5b).

## 3 Discussion

Here we report, that there were no longitudinal changes in resting-state functional connectivity (rsFC) between cortical motor areas despite substantial motor recovery over the same time period in a cohort of patients with subcortical stroke. In addition, at no stage of recovery were rsFC patterns different from healthy, age-matched controls.

Whenever results are negative, concerns will be raised about the power of the study (addressed below) and the biological validity of the method in general.

There have been more than 500 rs-fMRI studies of brain connectivity (Buckner *et al.*, 2013). Recent reports have described the close relationship between resting-state networks and structural connectivity assessed with other methods e.g. magnetoencephalography (van den Heuvel *et al.*, 2009; Brooks *et al.*, 2011). Most notably for our purposes, the sensitivity of rsFC to changes in experience-dependent neural plasticity appears to be quite high, as even short periods of training yield statistically significant changes of functional connectivity in small *n* studies in healthy subjects (Mawase *et al.*, 2017; Vahdat *et al.*, 2011). For example, Censor and colleagues (Censor *et al.*, 2014), in a comprehensive multimodal approach combining behavioral, brain stimulation, and rs-fMRI data, they demonstrated that changes in performance after training on a five-digit sequence task led to reliable changes in corticostriatal functional connectivity. When motor memory formation after training was disrupted using rTMS, changes in functional connectivity predicted the modification of memory recall on the next day.

Given such results, why were we not able to detect rsFC changes in the setting of stroke recovery? Injury ostensibly triggers functional reorganization, which arguably should be a more dramatic cause of connectivity change as it is associated with structural alterations, e.g. sprouting, and not just learning-related changes in pre-existing connections. There are two potential answers to this question, one is the possibility that the idea that changes in cortico-cortical connections promote motor recovery after stroke is ill-conceived, the second is that there are methodological limitations to rs-fMRI. We shall discuss both of these concerns.

A large number of animal studies, in rodents and non-human primates, have described numerous structural and physiological changes in cortical areas around and beyond the infarct core. These changes have collectively been called reorganization, but in only a small subset of cases have they been correlated with motor recovery, which suggests that most are likely just reactive (Carmichael, 2016). We reasoned that as spontaneous biological recovery is similar for cortical and subcortical strokes (Zarahn *et al.*, 2011) then recovery-related cortical reorganization, if not just reactive, should still occur in patients with isolated subcortical lesions. Indeed, we know that corticospinal integrity assessed with TMS is a good predictor of recovery in patients with subcortical stroke (Radlisnka *et al.*, 2010; Byblow et al. 2015), i.e., cortical output is required for recovery from subcortical stroke just like it is for cortical stroke. In addition, changes in cortical maps are seen not just with cortical lesions but with spinal and peripheral lesions as well (Florence *et al.*, 1998; Moxon *et al.*, 2014, Krakauer & Carmichael, 2017). Here, however, we found no evidence for systematic rsFC changes between cortical motor regions. In the light of these results, previously reported cortical connectivity changes could be reactive rather than reparative, e.g. confounded by the presence of a cortical lesion.

The question must now be asked why it was ever conjectured that changes in connections between cortical regions would enhance recovery from hemiparesis, which is caused by interruption of descending pathways out of a particular region(s). One could rephrase this to ask why would there be a “horizontal” solution to a “vertical” problem? This question is related to the increasing awareness of the questionable relevance of cortical map changes to recovery (Krakauer & Carmichael 2017), changes which have hitherto been taken as electrophysiological evidence for reorganization (Dancause & Nudo, 2011; Warraich & Kleim, 2010; Wittenberg, 2010). Overall, it is increasingly apparent both from recent and previous work in non-human primates and rodents that recovery after stroke relates to changes in the strengths of descending projections to the brainstem and spinal cord from individual motor cortical areas rather than to changes in the connections *between* them (Lin *et al.*, 2018; Starkey *et al.*, 2012; Wahl *et al.*, 2014; Zaaimi *et al.*, 2012). That said, it could be postulated that cortico-cortical drive, for example of premotor cortex onto primary motor cortex (M1) could facilitate remaining CST descending projections out of M1, as studies have shown such cortico-cortical facilitation in healthy non-human primates (Cerri *et al.*, 2003; Shimazu *et al.*, 2004). Consistent with what we found here, however, there is little evidence for this as a recovery mechanism after stroke in any animal.

While our results are congruent with similar observations in a smaller cohort (Nijboer *et al.*, 2017), they are seemingly contradicted by a recently published paper that reported results for resting-state changes in a similarly sized cohort of patients with subcortical stroke. In this study, Lee and colleagues obtained six connectivity measures between 40 supra- and infratentorial ROIs in 21 stroke patients measured at two time-points post-stroke (2 weeks and 3 months), and found differences in two of the measures. Specifically, they found lower overall strength in interhemispheric connectivity and higher network distance compared to healthy controls at 2 weeks, but neither measure changed at 3 months. Even if one overlooks the unmentioned comparisons problem and the fact that they had more variables (six measures, 40 ROIs) than subjects, their results showed no connectivity measure *changing* as the patients improved, which is consistent with our results.

Although we favor the view that the absence of connectivity change in our study is a true negative result both in terms of the power of the study and the biological validity of rsFC (van Meer *et al.*, 2010), an alternative explanation would relate to methodological limitations of rs-fMRI.

Methodological problems with e.g. regard to reproducibility of imaging analysis in general and rs-fMRI, in particular, have long been a topic of discussion (Baker, 2016; Ioannidis *et al.*, 2014; Macleod *et al.*, 2014). So far, there is no consensus about the optimal way to analyze rs-fMRI data, which poses a fundamental challenge regarding the generalizability and comparability of reported findings. In face of a low signal to noise ratio, missing consensus in analysis steps and statistical methods (promoting the risk of conscious and unconscious p-hacking; Nuzzo, 2015), and frequent absence of an *a priori* hypothesis (which can lead to so-called HARKing; Kerr, 1998), the imaging literature is especially vulnerable to false-positive or -negative results (Munafò *et al.*, 2017). For example, converging evidence highlights that the choice of different pre-processing strategies needs to be considered as an important confound in rs-fMRI (Cole *et al.*, 2010; He & Liu, 2012; Weissenbacher *et al.*, 2009).

We addressed this problem by providing measures of data reliability, comparing two different pre-processing procedures, and by reanalyzing our data set with regard to individual M1-M1 changes using a previously reported metric for resting-state imaging analysis (Golestani *et al.*, 2013). Here we provide, to the best of our knowledge, the most methodologically complete study to date of stroke recovery using rs-fMRI. Additionally, open science efforts including data sharing have been identified as a major tool to secure transparency and reproducibility of reported results, allowing for external validation of results, detection of mistakes, and generation of alternative interpretations (Nosek *et al.*, 2015). In an effort to increase the transparency and reproducibility of our results, the complete data set as well as the custom-written MATLAB and R scripts are made publicly available to invite further analysis.

### Conclusion

In the present study, we investigated longitudinal changes in functional connectivity after subcortical stroke. Despite substantial recovery from motor impairment over one year, we found no differences in functional connectivity between patients and controls, nor any changes over time. Assuming that rs-fMRI is an adequate method to capture connectivity changes between cortical regions after brain injury, the results presented here, provide reason to doubt that post-stroke cortical reorganization, conceived as changes in cortico-cortical connectivity, is the relevant mechanism for promoting motor recovery after stroke. We suggest instead that it is facilitation of residual cortical descending pathways that are likely to be more causally relevant. It is perhaps time for the field to change its emphasis from changes in “horizontal” connections to changes in “vertical” ones.

## 4 Materials and methods

The resting-state data set presented here was acquired from a natural history study investigating upper extremity recovery after stroke (Study of Motor Acute Recovery Time course after Stroke; SMARTS). As part of the study, a range of behavioral, physiological, and imaging measurements were obtained. Details of the behavioral characterization of the patients have been published elsewhere (Cortés *et al.*, 2017; Ejaz *et al.*, 2018; Xu *et al.*, 2017).

### 4.1. Patients

Since we were interested in cortical connectivity changes after stroke, in order to avoid confounding results due to cortical damage, only a subset of 19 patients with lesions restricted to subcortical areas was considered (6 females; mean age 59 ±12 years, 15 right-handed). Major inclusion criteria were: first-ever clinical apparent ischemic stroke, proven by a positive DWI lesion within the previous 2 weeks; unilateral upper extremity weakness (Medical Research Council muscle weakness scale <5); ability to give informed consent. Patients were excluded for one or more of the following reasons: initial impairment too mild (Fugl-Meyer score Upper Extremity >63/66), age ≤21 years, hemorrhagic stroke (Xu *et al.*, 2017). The selected patients had lesions either in the corona radiata, the internal capsule or in the cortico-spinal tract above the crossing in the pyramid. Demographics are described in Table S1; more detailed information about lesion distribution is shown in Figure S1.

Additionally, 11 healthy age-matched control participants (4 females; mean age 65 ±8 years; all right-handed), were tested at the same time-points.

The study was carried out in accordance with the Declaration of Helsinki and approved by the respective local ethics committee of the participating recruiting centers of SMARTS (Johns Hopkins University, USA, Columbia University, USA, University Hospital Zurich, Switzerland). All participants gave written informed consent.

### 4.2. Study design

Patients were enrolled in the study within the first two weeks after stroke and followed up over a one-year period at five time-points: acute stage: week 1-2 (10 ±4 days), W4: week 4-6 (37 ±8 days), W12: week 12-14 (95 ±10 days), W24: week 24-26 (187 ±12 days), and W52: week 52-54 (370 ±9 days). During each visit, the following clinical parameters were assessed: Fugl-Meyer score Upper Extremity (FM-UE, max. score 66, Fugl-Meyer *et al.*, 1975), Action Research Arm Test (ARAT, max. score 57, Yozbatiran *et al.*, 2008). Hand strength and individuation ability were measured using a custom-made hand-device (Xu *et al.*, 2017). The FM-UE and ARAT are widely used to assess motor deficits after stroke and can capture different aspects of recovery: higher FM-UE scores represent normal reflex activity, fewer muscular coactivations, coordination and higher joint mobility thought to be equal to “true” resolution of impairment; higher ARAT scores are achievable with compensatory strategies, thus correlating closer with activities of daily living. Measuring hand strength offers a third dimension of recovery that is only partially captured within the FM-UE and ARAT.

### 4.3. Image Acquisition

Participants were scanned with an 3T Achieva Philips system. Scans were obtained with a 32-channel head coil, using a two-dimensional echo-planar imaging sequence (TR=2.00s, 35 slices, 210 volumes/run, slice thickness 3mm, 1mm gap, in-plane resolution 3×3mm^2^). Each resting-state scan was 8min long. Participants were instructed to lie still and visually fixate on a central white cross displayed on a computer monitor. Structural images for atlas transformation and lesion definition were acquired with a T1-weighted anatomical scan (3D MPRAGE sequence, TR/TE=8/3.8ms, FOV 212×212mm, matrix 96×96, 60 slices, slice thickness 2.2mm). Finally, for each participant, a diffusion weighted imaging (DWI) image (TR=2.89s, 30 slices, 5mm slice thickness, 240×240mm FOV), was acquired to define lesion boundaries.

### 4.4. Imaging analysis

#### 4.4.1. Preprocessing of rs-fMRI time series

Rs-fMRI has a relatively low signal-to-noise ratio. Non-neuronal processes, such as sensor noise, head motion, cardiac phase, and breathing, account for a considerable part of the variance of the raw signal (Birn, 2012). It has been argued that markers for the reliability of the sampled rs-fMRI data are missing and that the choice of preprocessing steps is often not justified (Bennett & Miller, 2010; Zuo & Xing, 2014). We therefore conducted two different procedures for noise reduction and then compared split-half reliability for the whole connectivity pattern in controls to determine which steps provided higher reliability (see supplementary material).

#### 4.4.2. Lesion definition

Lesion boundaries were defined as an intensity increase of ≥30% on DWI images, and in a second step manually modified by a neuroradiologist and a neurologist using RoiEditor, see Figure S1 for averaged lesion distribution map.

#### 4.4.3. ROI definition

We chose five motor areas (S1=primary somatosensory cortex, M1=primary motor cortex, PMd=dorsal premotor cortex, PMv=ventral premotor cortex, SMA=supplementary motor area) as regions of interest that have been widely accepted as being associated with motor function and motor recovery (Miyai *et al.*, 1999, 2002; Rehme *et al.*, 2012). Individual T1-images were used to delineate pial-grey matter and grey matter-white matter boundaries using FreeSurfer software (Dale *et al.*, 1999). The cortical surfaces were aligned across participants based on the sulcal-depth and local curvature maps. Probabilistic cyto-architectonic maps (Fischl *et al.*, 2008) aligned to the group average surface were then used to define ROIs first on the individual surface, and then back-projected into the subject-native space.

The ROIs were defined as follows, M1: surface nodes with the highest probability for Brodmann area (BA) 4. To increase specificity for processes related to recovery of hand function, this ROI was limited to 2cm above and below the hand-knob (Yousry, 1997). S1: nodes in the hand-region in S1 were isolated using BA 3a, 3b, 1 and 2.2cm above and below the hand knob. PMd: nodes with highest probability in BA6, above middle frontal sulcus, but on the lateral surface of the hemisphere. PMv: nodes with the highest probability in BA6, above middle frontal sulcus. SMA: nodes with the highest probability in BA6 on the medial surface of the brain. This ROI therefore includes SMA and preSMA (Picard & Strick, 1996).

### 4.5. Functional connectivity analysis

For each ROI, the time series for all voxels within the ROI were extracted and averaged, resulting in a single BOLD time-course vector for each of the 10 ROIs across the two hemispheres (left-S1, left-M1, left-PMd, left-PMv, left-SMA, right-S1, right-M1, right-PMd, right-PMv, right-SMA). Pairwise correlations between averaged BOLD time-course vectors for the different ROIs were computed and Fisher-Z transformed to conform better to a normal distribution, resulting in a 10×10 matrix of connectivity weights (Figure 2). The matrix thus represents the connectivity weights between all possible ROIs for a patient: 10 intrahemispheric ROI pairs, each *within* the lesioned and non-lesioned hemispheres, respectively, and 25 interhemispheric ROI pairs *between* the lesioned and non-lesioned hemispheres (overall 45 connectivity weights for all ROI pairs). For the rest of this manuscript, this vectorized, Fisher-Z transformed correlation matrix will be referred to as the full connectivity pattern, while the corresponding intra- and interhemispheric subsets of the matrix will be referred to as the intrahemispheric non-lesioned (1×10 vector), intrahemispheric lesioned (1×10 vector), and interhemispheric connectivity patterns respectively (1×25 vector). These connectivity patterns were estimated independently for each session and patient. Connectivity patterns for controls were estimated similarly, with the exception that intrahemispheric connectivity patterns were averaged across both hemispheres.

### 4.6. Changes in connectivity patterns in the acute recovery period

In the early acute recovery period (week 1-2), stroke-related damage could alter connectivity patterns in patients in two distinct ways: 1) the connectivity pattern could remain the same but overall connection strengths might be increased or decreased, resulting in connectivity patterns in patients DC-shifted but otherwise identical to control patterns. This would indicate that a canonical pattern of connectivity between motor ROIs in healthy people is simply up or down-regulated post-stroke either due to maladaptation or compensation for damage. 2) stroke-related damage might alter connectivity weights among only a few select ROIs, e.g. either between ROIs within one hemisphere or across hemispheres. This would alter the shape of the connectivity patterns in patients in comparison to controls. Since we wanted to be sensitive to both kinds of connectivity pattern change, the appropriate statistical test would be a MANOVA between patient and control connectivity patterns. However, due to insufficient degrees of freedom in performing such an analysis (the number of connectivity weights exceeds the number of patients and controls), we instead opted for a permutation test with Euclidean distance as a measure of dissimilarity between patient and control connectivity patterns as it is sensitive to *shape and scaling* changes of connectivity patterns (for details see supplementary material).

While, on average, connectivity patterns for patients might not differ from controls in the acute recovery stage, individual patients might exhibit idiosyncratic connectivity patterns owing to the heterologous distribution of lesions locations in the cohort. Thus, acute stage changes in connectivity patterns might result in an increase in variability in within-group connectivity patterns. To determine whether this was the case in the acute stage, we computed the average Euclidean distances between each patient’s connectivity pattern and the patients’ mean connectivity pattern (acute P_variability). Similarly, we computed the average Euclidean distance between each individual control pattern and the controls’ mean connectivity pattern (acute C_variability). The differences between these two served as a measure of increased or decreased variability in the patients (P_variability-C_variability=Δvariability). We then repeated the permutation test (for details see supplementary material) to generate a null distribution of the difference in variability to test the significance of Δvariability.

### 4.7. Changes in connectivity patterns over time during recovery

Since patients in our cohort demonstrated substantial improvements of upper extremity deficits in the year after stroke (Figure 1), we were interested to see whether there were concomitant longitudinal changes in connectivity patterns. To determine this, we performed two separate but related analyses. First, we independently compared differences in patient connectivity patterns from the acute stage to all consecutive weeks (Δweek from acute to week 4, week 12, week 24, and week 52) to determine how far connectivity patterns diverged over the year from the pattern in the acute post-stroke stage. The same was done for control connectivity patterns to establish intersession reliability. Second, we compared patient’s connectivity patterns for all five measurement sessions against the control connectivity patterns to determine how patient patterns changed longitudinally in reference to controls (Δpattern for acute, week 4, week 12, week 24 and week 52). Both these analyses were performed using Euclidean distance and permutation testing in the same way as for estimating differences in connectivity patterns at the acute recovery stage.

To assess if individual idiosyncratic patterns might show a change over time that could underlie recovery, we analyzed individual connectivity pattern changes for a subgroup of patients with all time-points (10 patients) by comparing pattern variability in the acute stage against all other time-points (Δweek_variability for acute_week 4, acute_week 12, acute_week 24, and acute_week 52) and performing an ANOVA with the factor Weeks.

### 4.1. Alternative metrics to calculate functional connectivity

Because changes in functional connectivity between the two primary motor cortices have been reported more consistently than other connectivity changes after stroke, we also explicitly looked at changes of M1-M1 connectivity weights.

We additionally analyzed our dataset using a metric of functional connectivity that was proposed in the to-date largest longitudinal resting-state stroke study with cortical and subcortical lesion location, which reported changes of M1 interhemispheric connectivity. The metric has been called Relative connectivity (RelCon) and is claimed to have low sensitivity to the temporal signal-to-noise ratio and signal amplitude fluctuations while maintaining a high sensitivity to meaningful signal changes, therefore offering an advantage e.g. in the analysis of data sets acquired with different scanners (Golestani & Goodyear, 2011). RelCon looks at interhemispheric connectivity of M1 in relation to intrahemispheric connectivity of M1 (for details see supplementary material).

Based on the reported methods, we calculated the RelCon for interhemispheric SM1 connections in our dataset.

#### Statistical analysis

Changes of behavioral measures in patients over time were analyzed using a mixed-effects ANOVA, with Week (acute – W52) as a fixed factor, and Subject as a random factor. As approximately 11% of the sessions were missing, we used the lme4 toolbox in R (Bates *et al.*, 2015) to fit the unbalanced mixed-effects design. Rather than F-values, statistical tests for main effects and interactions are reported using a χ^2^ approximation. Behavioral measures of patients and controls at the acute stage were compared with a two-tailed t-test.

Intrasession reliability was analyzed by computing split-half correlations (Pearson’s correlation) for each single week and individual patient/control, as well as looking at the averaged split-half correlation for all weeks together. Reliability between groups was compared using a mixed-effects ANOVA, with Group (patients vs. controls) and Week (acute – W52) as fixed, and Subject as a random factor. This was done for all connections, as well as subsets only including interhemispheric, intrahemispheric lesioned or non-lesioned ROIs.

Changes of interhemispheric M1-M1 connectivity weights over time between patients and controls were analyzed using a mixed-effects ANOVA, with Group (patients vs. controls) and Week (acute – W52) as fixed, and Subject as a random factor, alternative metrics reported in Golestani et al. were analyzed in the same way.

Results were considered significant at p<0.05. Means values are reported ± standard deviation unless stated otherwise.

#### Data availability

The complete data set will be openly available in a public repository upon publication. All analysis was performed using built-in and custom-written MATLAB and R scripts that will be made publicly available upon publication.

## Acknowledgement

We like to thank Susumu Mori & Andreia Faria from the Department of Neuroradiology, Johns Hopkins University for their support regarding imaging analysis.

## 7. Supplemental Material

### Methods

#### Permutation test and Bootstrapping

To perform a permutation test, we first identified patients and controls that had estimates of connectivity patterns within the first two weeks after stroke. We estimated Δpattern as the Euclidean distance between the average connectivity pattern for patients and the average connectivity pattern for controls. We then shuffled group assignment labels for connectivity patterns 10,000 times, randomly assigning connectivity patterns to “controls” or “patients”. From the shuffled data, we again calculated the Euclidean distance between the average connectivity pattern for patients and controls based on this new assignment. By repeatedly shuffling and computing Euclidean distances, we obtained an estimate of the empirical null distribution of Δpattern – e.g. the expected distribution if there was no real difference between the two groups. The measured Δpattern was then compared against this null distribution, and the relative proportion of simulations that showed a larger distance was used as a p-value - the probability that the distance between the mean control and patient pattern would be equal or larger than the measured distance by pure chance. This analysis was carried out independently for the full, intrahemispheric lesioned, intrahemispheric non-lesioned, and interhemispheric connectivity patterns.

#### RelCon

To calculate the interhemispheric RelCon for ipsilesional and contralesional sensorimotor cortex (SM1) the correlation between time-series of all possible pairs of voxels is calculated (all voxels SM1_ipsilesional-contralesional_). The average of the interhemispheric connectivity for SM1_ipsilesional-contralesional_ is then calculated relative to the within connectivity of the ipsilesional SM1 (divided by the average correlation of all voxel within SM1_ipsilesional_).

This metric was tested on different real and simulated data sets and showed superior results compared to other *absolute* connectivity measures (absolute meaning connectivity measures that do not relate interhemispheric ROI-to-ROI connectivity weights to the average within correlation of the ipsilesional ROI itself).

#### Results

**Table S1:** Patient demographics and overall session count. First FM-UE = first recorded Fugl-Meyer score Upper Extremity, first ARAT = first recorded Arm Research Action Test.

| ID | age | gender | handedness | lesion side | first FM-UE | first ARAT | session |
| --- | --- | --- | --- | --- | --- | --- | --- |
| 2310 | 57 | m | right | left | 58 | 56 | 5 |
| 2365 | 53 | f | right | right | 0 | 57 | 4 |
| 2395 | 65 | m | right | right | 30 | 21 | 4 |
| 2450 | 66 | m | right | right | 66 | 56 | 3 |
| 2531 | 66 | f | right | right | 60 | 55 | 5 |
| 2565 | 71 | m | right | right | 4 | 0 | 3 |
| 2652 | 46 | m | left | left | 4 | 0 | 4 |
| 2654 | 46 | m | right | right | 49 | 52 | 5 |
| 2663 | 67 | f | right | left | 16 | 2 | 4 |
| 2789 | 56 | m | right | right | 64 | 57 | 4 |
| 2925 | 59 | f | right | left | 60 | 57 | 5 |
| 3176 | 64 | m | left | right | 63 | 57 | 4 |
| 3239 | 74 | m | left | left | 5 | 0 | 5 |
| 3240 | 80 | f | right | left | 9 | 56 | 5 |
| 3241 | 64 | f | right | right | 58 | 39 | 5 |
| 3243 | 22 | m | right | left | 63 | 56 | 5 |
| 3246 | 53 | m | left | left | 30 | 39 | 5 |
| 3247 | 54 | m | right | right | 59 | 57 | 5 |
| 3248 | 58 | m | right | right | 61 | 56 | 4 |

**Figure S1:**
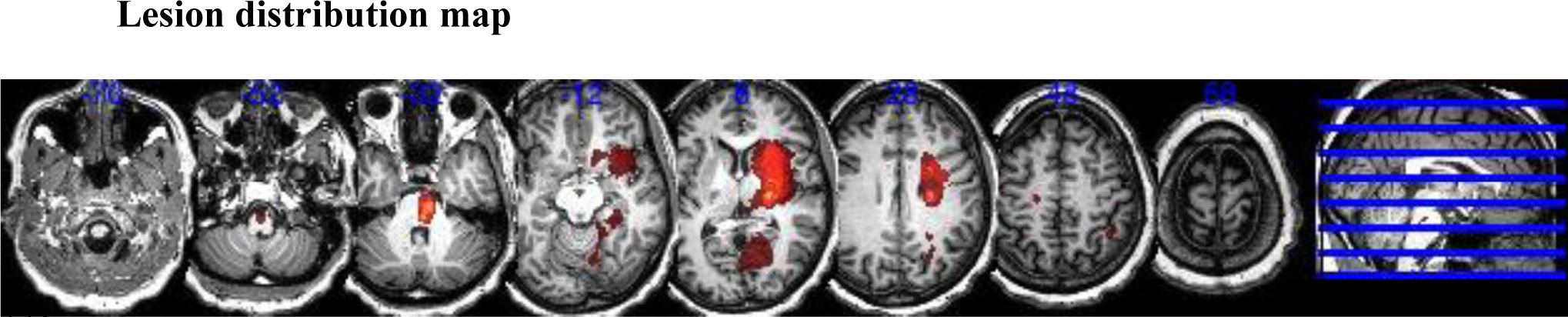
Lesion distribution of patients (N = 19). Averaged lesion distribution mapped to MNI space with lesion flipped to one hemisphere.

#### 2.6 Data reliability and Preprocessing comparison

To estimate the reliability of our measurements within sessions, connectivity patterns were computed as described above for the first 100 volumes and the second 100 volumes independently and correlated with each other to calculate split-half reliabilities.

**Figure S2:**
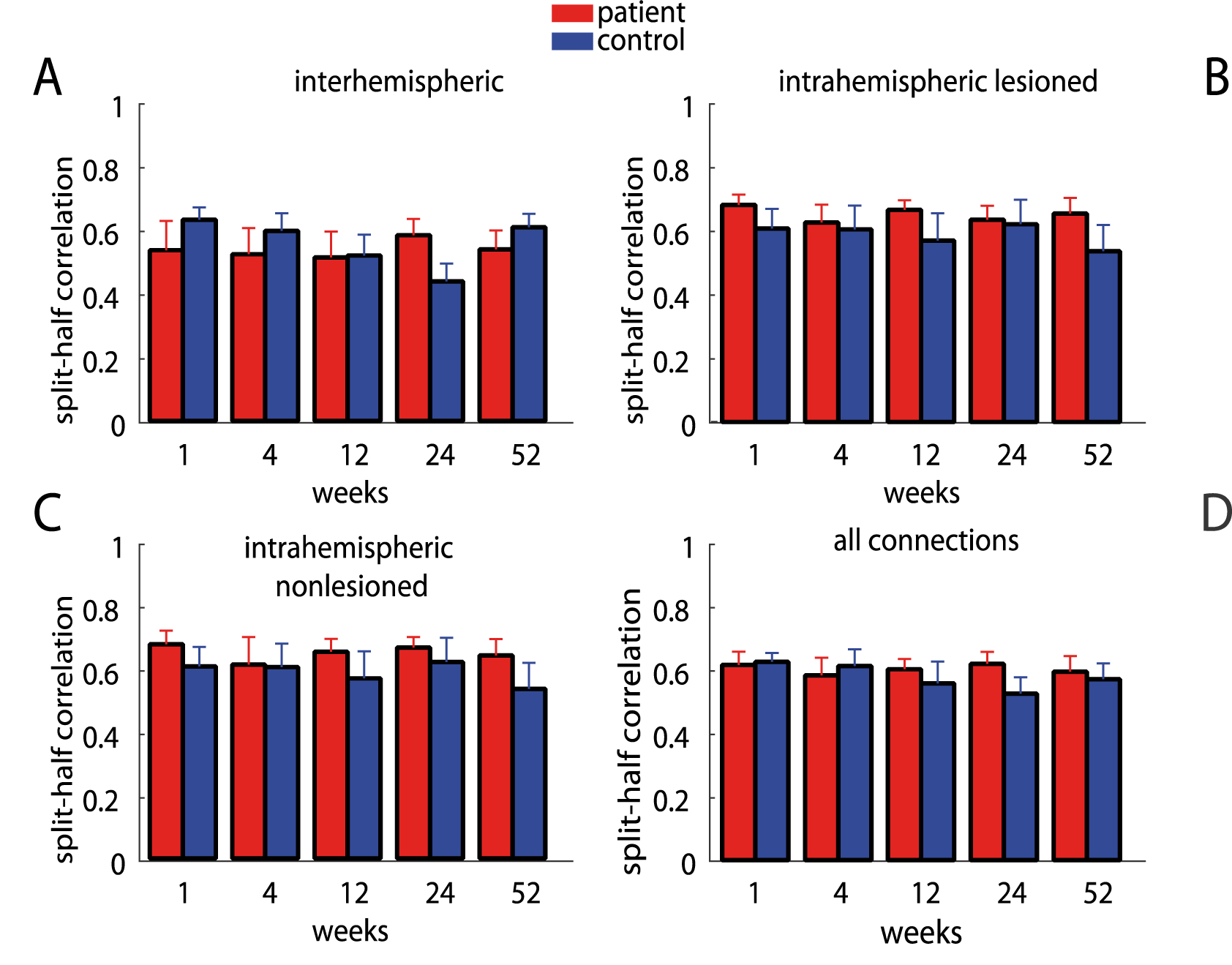
Intrasession split-half reliability for patients and controls at each week. For each individual patient or control participant, the BOLD time series of the first 100 volumes of the scan were correlated with the last 100 volumes. Each Panel shows the results for a different condition. Patients always had slightly higher intrasession reliability than although this difference was not significant and was possibly driven by outlier in the control group.

As seen for overall connectivity, intra- and interhemispheric split-half reliabilities were highly reliable for controls and patients (controls: *intrahemispheric*: r = 0.67 (95% Confidence Interval, 0.61-0.74), *interhemispheric*: r = 0.64 (CI 0.59-0.69), patients: *intrahemispheric lesioned*: r = 0.78 (CI 0.75-0.81), *non-lesioned*: r = 0.77 (CI 0.73-0.82), *interhemispheric*: r = 0.69 (CI 0.66-0.74), see Figure S1 for split-half reliability of each week). Split-half reliability was not different between groups for *interhemispheric*: (χ^2^(1) = 0.0239, p = 0.8771) and *intrahemispheric non-lesioned* connections: (χ^2^(1) = 3.5634, p = 0.0591), but was different for *intrahemispheric lesioned* connections (χ^2^(1) = 4.2337, p = 0.0396).

The reliability measurement also allowed us to compared two different pre-processing procedures:

Preprocessing procedure (P1): We removed the first 10 volumes of the functional data, then performed correction for the timing of slice acquisition, motion correction, brain extraction, linear trend removal, and temporal filtering (band pass, 0.01-0.08 Hz) using FSL (FMRIB Software Library (FSL), Oxford University, Oxford, UK). Our analysis was carried out in the native space, and no spatial smoothing was applied. Linear regression was used to remove signal correlated with the global mean signal, and the average time series in the cerebral white matter and cerebrospinal fluid (Fox *et al.,* 2006).

Preprocessing procedure (P2): Here, we used an independent component analysis (ICA) approach using FSL MELODIC for artifact reduction (Smith *et al.*, 2004). Again, we removed the first 10 volumes of the functional data. We applied motion correction and brain extraction. Probabilistic independent component analysis was conducted to denoise individual data by removing components such as head motion, scanner artifacts, and physiological noise. Noise components were classified using FMRIB’s ICA-based Xnoiseifier (Salimi-Khorshidi *et al.*, 2014), which attempts to auto-classify ICA components into “good” vs. “bad” components. The “bad” components were then removed from the functional data.

To determine which procedure would provide a more stable result, we calculated the split-half reliability of the ROI-ROI connectivity weights for the whole connectivity pattern over time in controls only.

Both procedures lead to good intrasession reliability on average (P1 = 0.64, CI 0.60–0.66; P2 = 0.62, CI 0.57–0.66) but showed no significant difference (χ^2^(1) = 1.231, p = 0.267), while no consistent change over time was found for either procedure by itself (P1: χ^2^(4) = 2.834, p = 0.684; P2: χ^2^(4) = 3.007, p = 0.557). Because of the nominal higher intrasession reliability we conducted all subsequent analyses after noise correction using the P1 procedure.

As for overall connectivity, the intersession reliability for controls showed no significant change over time for intra- or intrahemispheric (*intrahemispheric lesioned:* Δweek acute_W4 = 0.36, CI 0.217–1.308; acute_W12 = 0.323, CI 0.225–1.119; acute_W24 = 0.567, CI 0.259–1.333; acute_W52 = 0.527, CI 0.228–1.118; *intrahemispheric non-lesioned:* Δweek acute_W4 = 0.36, CI 0.216–1.286; Δweek = 0.323, CI 0.221–1.121; acute_W24 = 0.567, CI 0.26–1.331; acute_W52 = 0.527, CI 0.23–1.136; *interhemispheric:*, Δweek acute_W4 = 0.669, CI 0.445–2.062; acute_W12 = 0.516, CI 0.419–1.876; acute_W24 = 0.721, CI 0.519–2.127; acute_W52 = 0.751, CI 0.45–1.991).

#### 7.2.1 Homo- versus Heterologous ROI connectivity

The physiological plausibility of the recorded BOLD signal fluctuations was further examined by comparing functional connectivity of homologous versus heterologous interhemispheric ROI-ROI connectivity weights using a linear mixed-model with participants as random factor and type (homo- or heterologous), week and group (control vs patients) as fixed factors, supplemental results Figure S3.

#### Homo- versus Heterologous ROI connectivity

We examined the differences in connectivity between homo- and heterologous ROI connection. Homologous connectivity was significantly higher than connections between heterologous ROIs (χ^2^(1) = 108.38, p<0.001) and this effect showed no changes over time χ^2^(4) = 5.8993, p = 0.207), Figure S1. Furthermore, we found no differences for this effect between patients and controls (type*group χ^2^(1) = 2.2701, p = 0.132 and type*week*group χ^2^(1) = 2.2187, p = 0.136), Figure S3.

**Figure S3:**
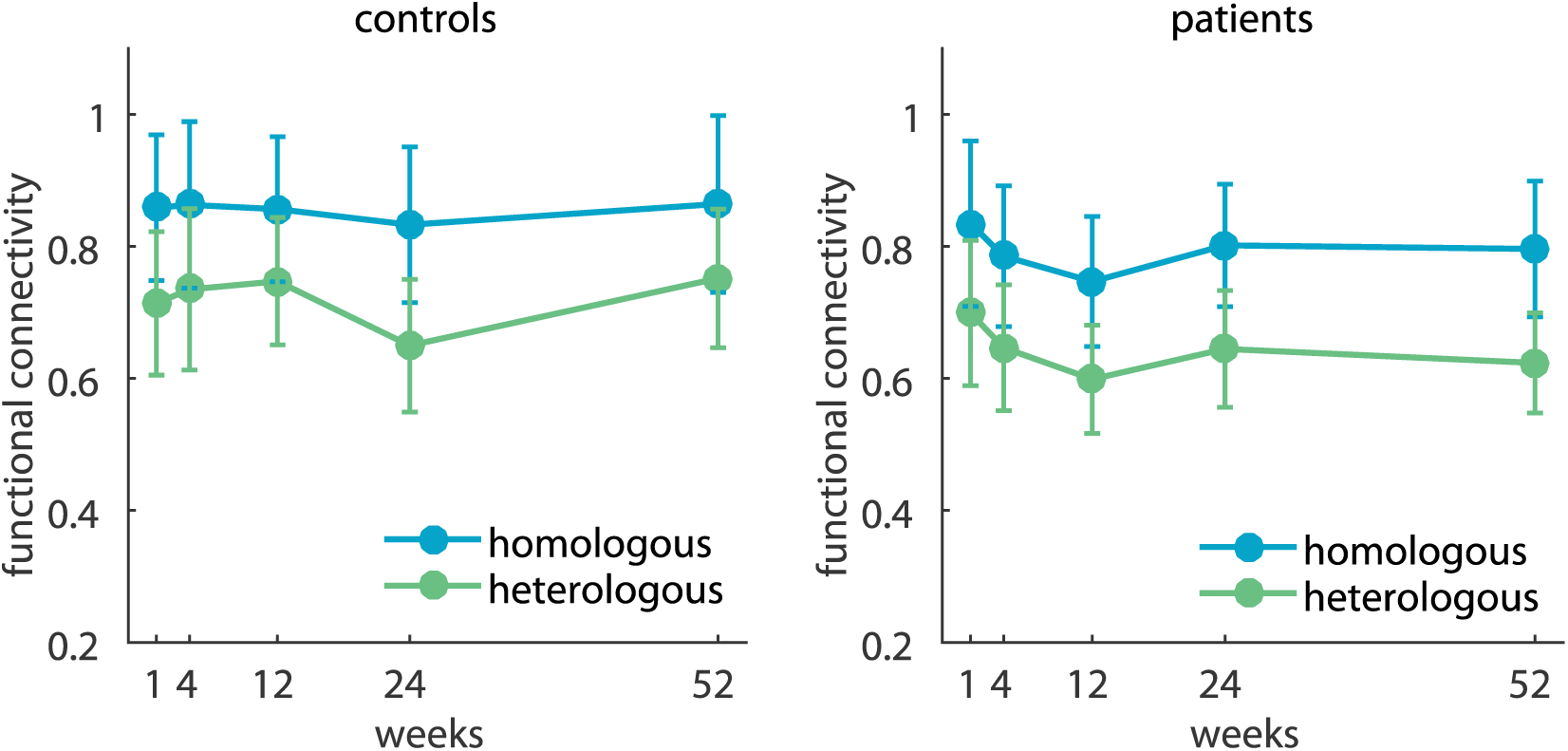
Average connectivity of interhemispheric homologous versus heterologous ROI-ROI connectivity weights. Homologous regions (e.g. M1-M1, S1-S1, blue line) were higher correlated than heterologous regions (e.g. M1-PmV, PmV-S1, green line). This did not change over the course of a year and no systematic difference was found between both groups (patients right panel, control left panel).

#### 7.2.2 Correlations between patient and control connectivity patterns

Connectivity patterns for patients and controls were highly correlated in the early period after stroke as well as at all subsequent measured time-points over the year (*all connections*: W1: R = 0.69, p = 0.0002; W4: R = 0.74, p<0.0001; W12: R = 0.76, p<0.0001; W24: R = 0.87, p = 0.0001; W52: R = 0.80, p<0.0001; *interhemispheric*: W1: R = 0.67, p = 0.0002; W4: R = 0.71, p<0.0001; W12: R = 0.73, p<0.0001; W24: R = 0.87, p<0.0001; W52: R = 0.81, p<0.0001; *intrahemispheric lesioned*: W1: R = 0.95, p<0.0001; W4: R = 0.96, p = 0.0088; W12: R = 0.89, p = 0.0006; W24: R = 0.96, p = 0.0129; W52: R = 0.96, p<0.0001; *intrahemispheric non-lesioned*: W1: R = 0.86, p = 0.0014; W4: R = 0.98, p = 0.0089; W12: R = 0.94, p<0.0001; W24: R = 0.94, p = 0.0001; W52: R = 0.89, p = 0.0006; Figure S4).

**Figure S4:**
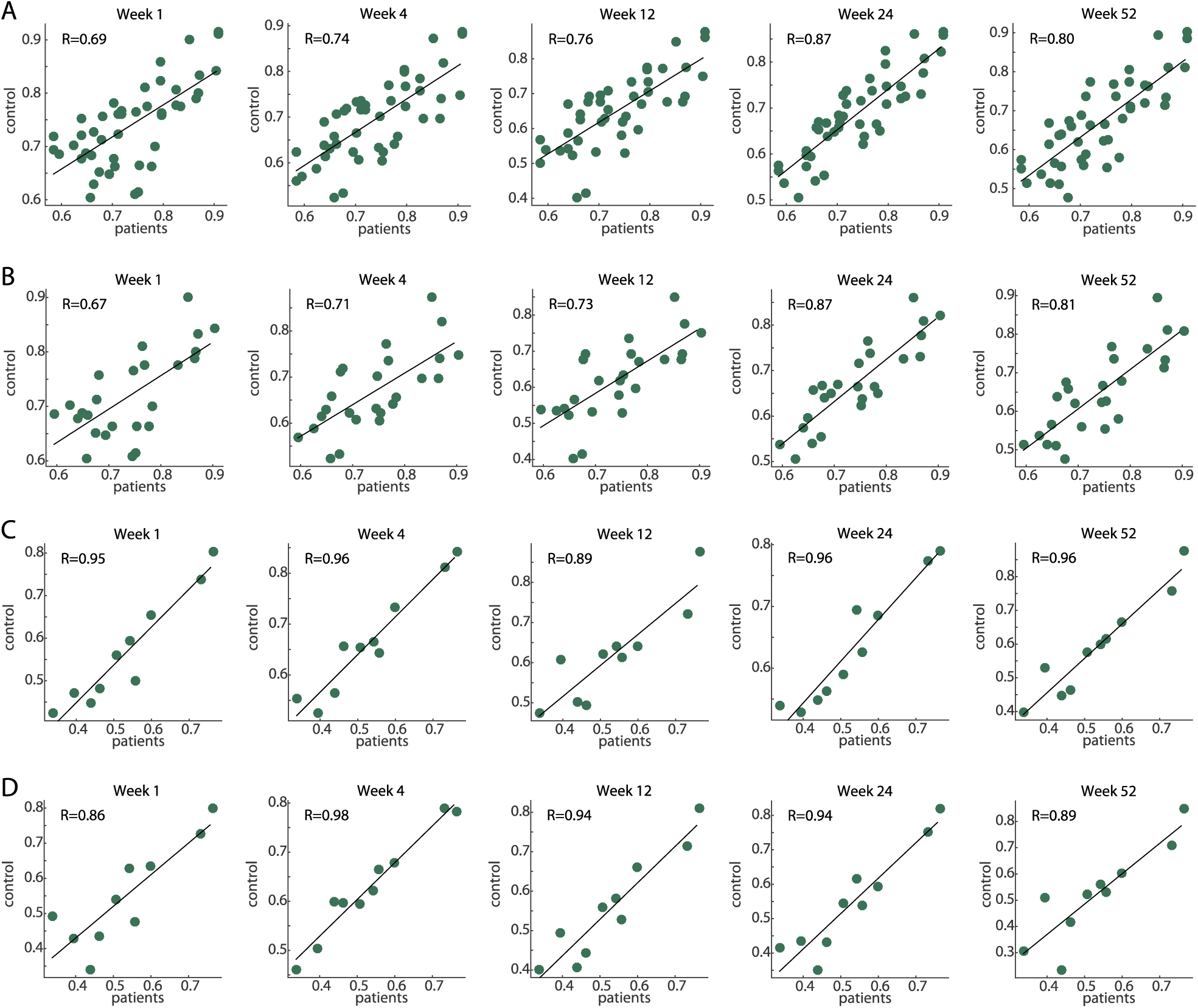
Correlations of connectivity patterns for patients (x-axis) and controls (y-axis) at each week. A) all connections, B) interhemispheric, C) intrahemispheric lesioned, D) intrahemispheric non-lesioned connectivity patterns.

